# Social touch observation in adults with autism: intact neural representations of affective meaning but lack of embodied resonance

**DOI:** 10.1101/512327

**Authors:** Haemy Lee Masson, Ineke Pillet, Steffie Amelynck, Stien Van De Plas, Michelle Hendriks, Hans Op de Beeck, Bart Boets

**Author notes:** H.O. and B.B. contributed equally to this work. To whom correspondence should be addressed. H. Lee Masson Department of Brain and Cognition, KU Leuven Tiensestraat 102, box 3714, Leuven, 3000, Belgium.

## Abstract

Humans can easily grasp the affective meaning of touch when observing social interactions. Several neural systems support this ability, including theory of mind (ToM) and somatosensory resonance systems, but it is unclear how these systems are affected in autism spectrum disorder (ASD). Individuals with ASD are characterized by impairments in social interaction and the use of (non)verbal communication such as social and reciprocal touch. The present study applies an ecologically valid stimulus set and multivoxel pattern fMRI neuroimaging to pinpoint atypicalities in the neural circuitry underlying socio-affective touch observation in adults with ASD as compared to matched neurotypical controls. The MVPA results reveal that the affective meaning of touch is well represented in the temporoparietal junction, a core ToM mentalizing area, in both groups. Conversely, only the neurotypical group hosts affective touch representations in the somatosensory cortex, not the ASD group, yielding a significant group difference. Lastly, individuals with a more positive attitude towards receiving, witnessing, and providing social touch and with a higher score on social responsivity, show more differentiated representations of the affective meaning of touch in these somatosensory areas. Together, our findings imply that individuals with ASD are able to cognitively represent the affective meaning of touch, but they lack the spontaneous embodied somatosensory resonance when observing social touch communications. Individual differences in this diminished resonance appear to be related to social touch avoidance and quantitative autism traits.

**Significance Statement:** Autism is characterized by socio-communicative impairments, including abnormal processing of interpersonal touch. Little is known about the neural basis of atypicalities in social touch processing in autism. Here, adults with and without autism watched video clips displaying social touch interactions and judged the affective valence of the touch. Subsequently, they underwent functional magnetic resonance imaging while watching the same videos. Brain activity patterns demonstrate that adults with autism show intact cognitive understanding (i.e. “knowing”) of observed socio-affective touch experiences but lack of embodied emotional resonance (i.e. “feeling”). This lack of emotional resonance is linked to social touch avoidance and quantitative autism traits. These findings highlight that the depth of experiencing the state of others is shallower in people with autism.

## Introduction

Interpersonal touch such as a hug and a slap is a potent non-verbal communicative tool for expressing one’s emotions and intentions [1], [2], thus a proper understanding is crucial for social functioning. Humans can extract a vast amount of information, including other’s affective states, when merely watching a touch interaction [3], [4]. Identifying other’s emotions from these social cues involves a sophisticated neural circuitry, including the extended visual system, the limbic system [5], and regions implicated in social cognition [6].

Pertaining to social cognition, two complementary theoretical frameworks – along with their associated neural modules – have targeted the processing of emotional body language. The first aligns with the more cognitively oriented Theory of Mind account (ToM; both the modular-theory and theory-theory), and postulates that humans are able to infer other’s mental states (i.e. emotions, intentions, and beliefs) by means of meta-perspective reasoning [7]–[9]. This ToM system has been depicted as a relatively effortfull, controlled and cognitively demanding form of social cognition [10], implicating the bilateral temporoparietal junction (TPJ) [11], [12]. The second account originates from the embodied simulation literature and aligns with the mirror neuron mechanism theory, and posits that individuals implicitly infer other people’s emotional states from social cues by automatically re-enacting pre-acquired sensory experiences [13], [14].

While this second line of research initially focused on the observation of fairly simple motor activities, implicating the premotor cortex and inferior parietal areas [15], [16], more recent studies have started investigating the observation of simple touch [5], [17], [18] and more complex interpersonal touch [19]. Concerning touch observation, accumulating evidence suggests that activated brain regions go beyond the visual cortex and include somatosensory regions involved in the processing of self-experienced touch [17], [20]–[22]. This direct mapping of other’s bodily experiences to the self may aid in simulating and empathizing with others’ emotions (e.g., the pain we feel when we observe another person being injected with a needle). Accordingly, the level of activation in the somatosensory system during touch observation has been associated with interindividual differences in empathy [23]–[26].

Although many studies have affirmed the presence of interindividual differences in social cognition, the behavioral and neural mechanisms of social touch perception have not been thoroughly investigated in neuropathological populations. Among the most relevant in this context is autism spectrum disorder (ASD), a hereditary neurodevelopmental disorder that is characterized by impairments in social interaction and communication and the presence of restricted, repetitive and stereotyped patterns of behavior [27]. ASD is often accompanied by an aversion to social touch [28], [29]. Using a limited range of touch stimuli, previous studies have shown that individuals with ASD frequently struggle with both receiving and offering touch [28]–[30], display reduced empathic resonance to painful touch observation [31], and show diminished neural activity in social brain regions in response to pleasant, gentle touch [32].

While difficulty in interpreting other people`s emotions from non-verbal social cues such as facial [33], [34] and bodily expressions [35], [36] is one of the diagnostic criteria of ASD, the empirical evidence in experimental studies is mixed [37], [38]. At a theoretical level, the socio-communicative impairments of individuals with ASD have often been attributed to impaired ToM abilities [7], [39], as well as to deficits in spontaneous social resonance [40]–[42].

Initial studies showed impaired or delayed development of ToM abilities in ASD, as evidenced by deficits in perspective taking, false belief processing and emotion recognition [43]. Likewise, at a neural level, individuals with ASD showed attenuated brain activity in the TPJ during various socio-cognitive tasks targeting ToM [44]–[46] (but see [47], [48]). On the other hand, it has been gradually recognized that many individuals with ASD, especially the higher functioning and verbally gifted ones, are able to pass these ToM tasks by means of compensatory sensory strategies and rule-based reasoning [49] despite substantial impairments in spontaneous social communication and interaction in daily life. This is where the embodied simulation account comes into play. According to this account, social impairments in ASD may result from a disability to simulate observed actions and internal states of others via personal sensory and emotional representations [50]. This account is supported by reduced brain activation in the mirror neuron system (MNS) of individuals with ASD during a variety of tasks requiring simulation [40], [41], [51], [52] (but see [53] for an argument against the broken mirror theory of ASD). Yet, this account has mainly been investigated with rudimentary action observation paradigms, and should be studied using more socially relevant and ecologically valid interactive paradigms.

The current study aims at understanding socio-affective touch processing in adults with ASD, both at the behavioral and neural level, using a stimulus set with complex and ecologically valid videos of social touch interactions. We particularly aim at unraveling whether neural representations of socio-affective touch observation are represented in a cognitive rule-based manner or based on embodied somatosensory resonance. We also investigate to what extent individual differences in socio-affective representations in brain regions relate to the presence of autism symptoms and touch aversion.

## Results

### Affective responses to social and non-social touch videos

Overall, both groups perceived the affective meaning of the touch videos as expected. More specifically, positive touch videos were rated as pleasant, negative touch as negative, and non-social touch as neutral. Concerning arousal ratings, both groups perceived social touch as exciting and non-social touch as calm. Fig. 1. shows data points of all individual participants for valence (a) and arousal (b) ratings. In addition, within- and between-subjects reliability tests revealed that participants were consistent in their ratings between the two sessions and were consistent with each other within each group. Summary statistics, statistical inference and an additional figure can be found in Supplemental Information (SI, Fig. S1).

**Fig. 1.**
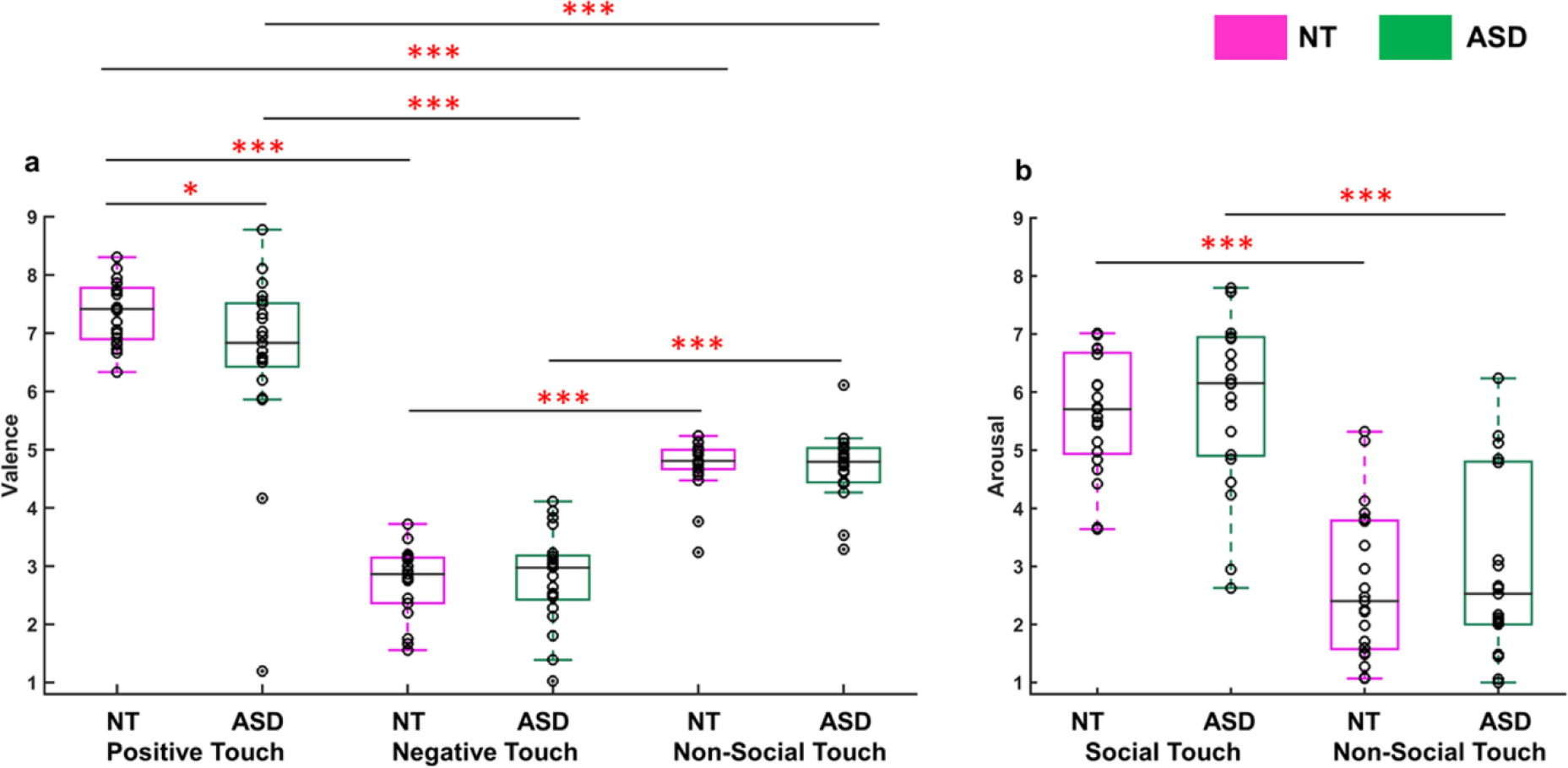
Affective responses to social (positive and negative) and non-social touch stimuli. The boxplots show each group’s valence (a) and arousal ratings (b) across conditions. The black lines inside of each box indicate the group medians. The bottom and top border edges of the boxes illustrate the 25^th^ and 75^th^ percentiles respectively; the whiskers illustrate the range of the individual ratings within 99.3 % coverage. The data points from individual participants’ ratings are additionally marked as black circles. The black dots outside of the boxes indicate the outliers. The single and three red asterisks indicate the statistical significance at p < 0.05 and p < 0.001 respectively. The x-axis indicates the name of conditions (e.g., positive touch) and the group (NT and ASD).

Regarding the difference between groups, a Mann-Whitney U test revealed no group difference in valence ratings of negative (z = −0.21, p = 0.83) and non-social touch videos (z = 0.42, p = 0.68). On the contrary, we observed a significant (albeit small) difference in the rated valence of the positive videos between the two groups (z = 1.99, p = 0.046), indicating that participants with ASD perceived positive social touch, such as a hug, as relatively less pleasant. Although outlying data points were observed in the valence ratings (Fig. 1a), the Mann-Whitney U test can robustly handle this as it is based upon medians rather than means. Both groups did not differ in their arousal ratings of social (z = −0.97, p = 0.33) and non-social touch videos (z = −0.40, p = 0.69).

### Social touch preference and its association with quantitative autism traits

Individuals with ASD (M = 56.8, SD =13.2) showed a less positive appreciation towards giving, receiving, and witnessing social touch in daily life (STQ: t (40) = 3.21 p = 0.003), as compared to the NT group (M = 69.2, SD = 9.6). As expected, individuals with ASD show a higher number of autism traits, as compared to NT individuals (SRS-A: t (40) = −3.72 p < 0.001). This group difference was significant on each of the social deficit subscales: social awareness (t (40) = −3.18, p = 0.003); social communication (t (40) = −3.33, p = 0.002); social motivation (t (40) = −2.88, p = 0.006).

The correlational analysis revealed a negative linear association between individual differences in social touch preference and the number of autism traits experienced by an individual (all participants r = −0.62, p < 0.001; NT group r = −0.55, p = 0.009; ASD group r = −0.48, p = 0.03, Fig 2). Similar associations were present for each of the social deficit subscales (SI, Table S1). Together, our results confirm that individuals with ASD present social impairments and exhibit a higher degree of social touch avoidance. Furthermore, the participants avoiding social touch seem to exhibit stronger social impairment, implying a tight link between social touch aversion and autism symptom severity.

**Fig. 2.**
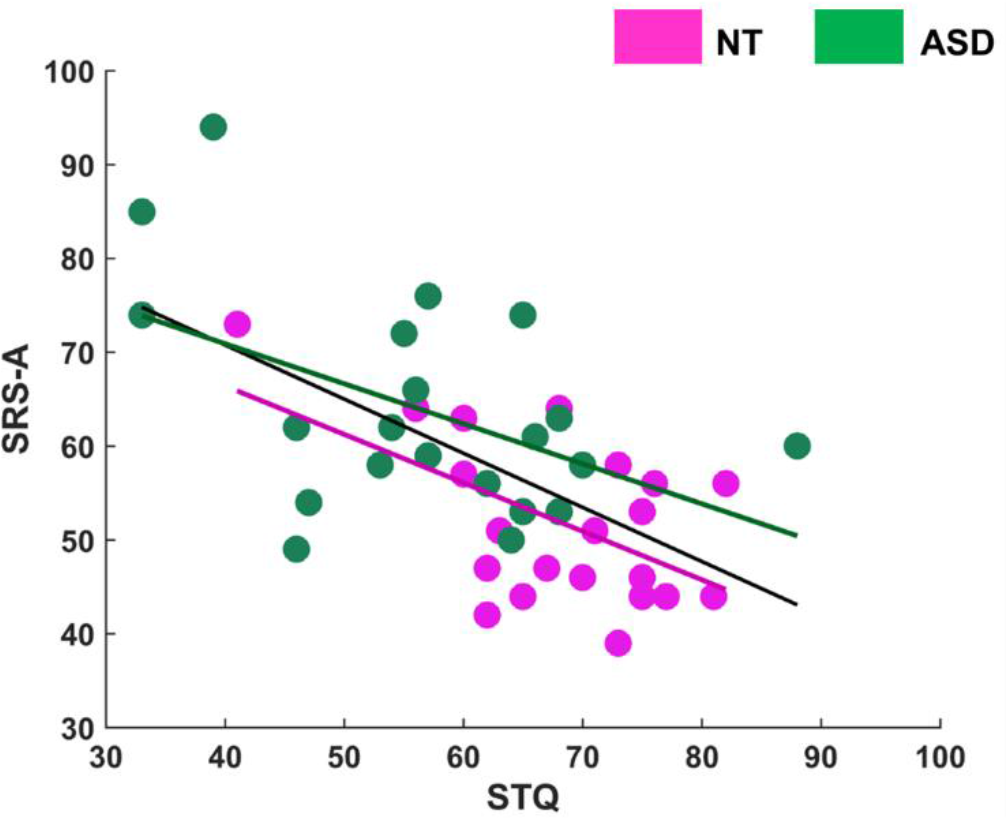
Social touch preference and its association with SRS-A scores. The figure illustrates the relationship between STQ and SRS-A scores. Individual data points of ASD and NT groups are marked as green and pink circles, respectively. The green, pink and black trend lines respectively indicate the negative association between the two variables in the ASD group, the NT group, and across all participants.

### Univariate neural responses to observed and felt touch

Two-sample t-tests revealed no significant group difference in neural responses for the contrast of social vs. non-social touch videos and vice versa (not at p fwe < 0.05, and not even with the lenient threshold p _uncorrected_ < 0.001). Mean group effects [0.5 0.5] of social vs. non-social touch contrasts are shown in Fig. 3 (SI, Table S2 for detailed information such as MNI coordinates of peak activity). Similarly, no significant group difference in neural responses for felt touch was found (not at p fwe < 0.05 and not at p _uncorrected_ < 0.001) (SI, Fig. S2 and Table S3).

**Fig. 3.**
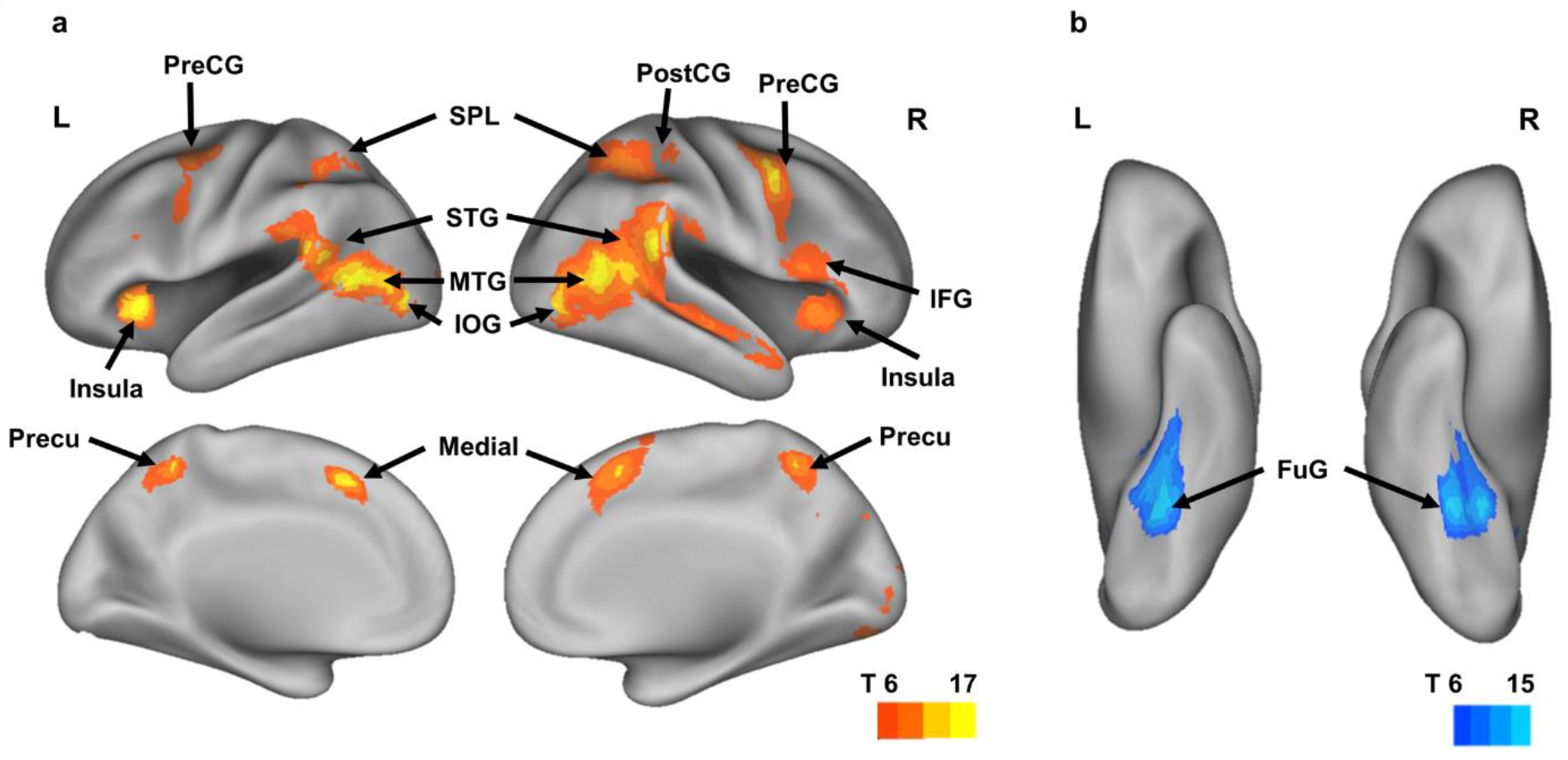
Brain areas involved in social vs. non-social touch observation. Figure (a) shows the significant differences (univariate random-effects between-subject analysis) in the contrast of social touch minus non-social touch while figure (b) shows the reverse contrast of non-social touch minus social touch (p fwe < 0.05, k= 60). The two groups did not differ in univariate activity. We mapped both contrast results on inflated cortices with PALS atlas using CARET software [70], [71]. L = left hemisphere, R = right hemisphere, PreCG = precentral gyrus, PostCG = postcentral gyrus, SPL = superior parietal lobe, STG = superior temporal gyrus, MTG = middle temporal gyrus, IOG = inferior occipital gyrus, Medial = medial prefrontal cortex, Precu = precuneus, FuG = fusiform gyrus, T = t-values

### Neural representations underlying observed social versus non-social touch processing

In line with earlier research by others [54] and ourselves [19], we expect that on top of this univariate selectivity there will also be high multi-voxel selectivity for the distinction between social and nonsocial touch videos. The multiple regression analysis confirmed that almost every implicated ROI represents the distinction between social vs. non-social touch scenes, even after controlling for the effects of all the other regressor variables (e.g., low-level visual features and motor response). Fig. 4a displays the main results and more details can be found in SI. Importantly, no significant group differences in neural selectivity for this distinction were found in the core ToM area (TPJ (t (40) = −0.67, p = 0.50)) and the somatosensory areas (BA3 z = 0.93, p = 0.35; BA1 z = −0.05, p = 0.96; BA2 z = −0.31, p = 0.76), indicating well-preserved neural selectivity in the ASD group for the social vs. non-social aspects shown in touch actions of others.

Note that these and all the following multi-voxel analyses require a reproducible signal, and between-group comparisons are easier to interpret if the reliability is comparable between the two groups. To assess whether there may be group differences in the reliability of the neural data in the TPJ and the somatosensory cortex, we calculated values of the leave-1 subject-out correlations within each group (correlating the neural data of one subject with the group averaged neural data after excluding this subject). Our results demonstrated that there was no group difference in the reliability of neural patterns in the four ROIs that are central to the tested hypotheses (BA3 t (40) = −1.75, p = 0.09; BA1 t (40) = 0.40, p = 0.69; BA2 t (40) = −0.24, p = 0.81; TPJ t (40) = 1.40, p = 0.17), (SI, Fig. S3).

In sum, our results suggest that the brains of individuals with and without ASD can equally distinguish whether other person’s touch actions comprise social interactions or not, and this rudimental social processing is implemented across multiple brain areas including visual, somatosensory and social regions.

**Fig. 4.**
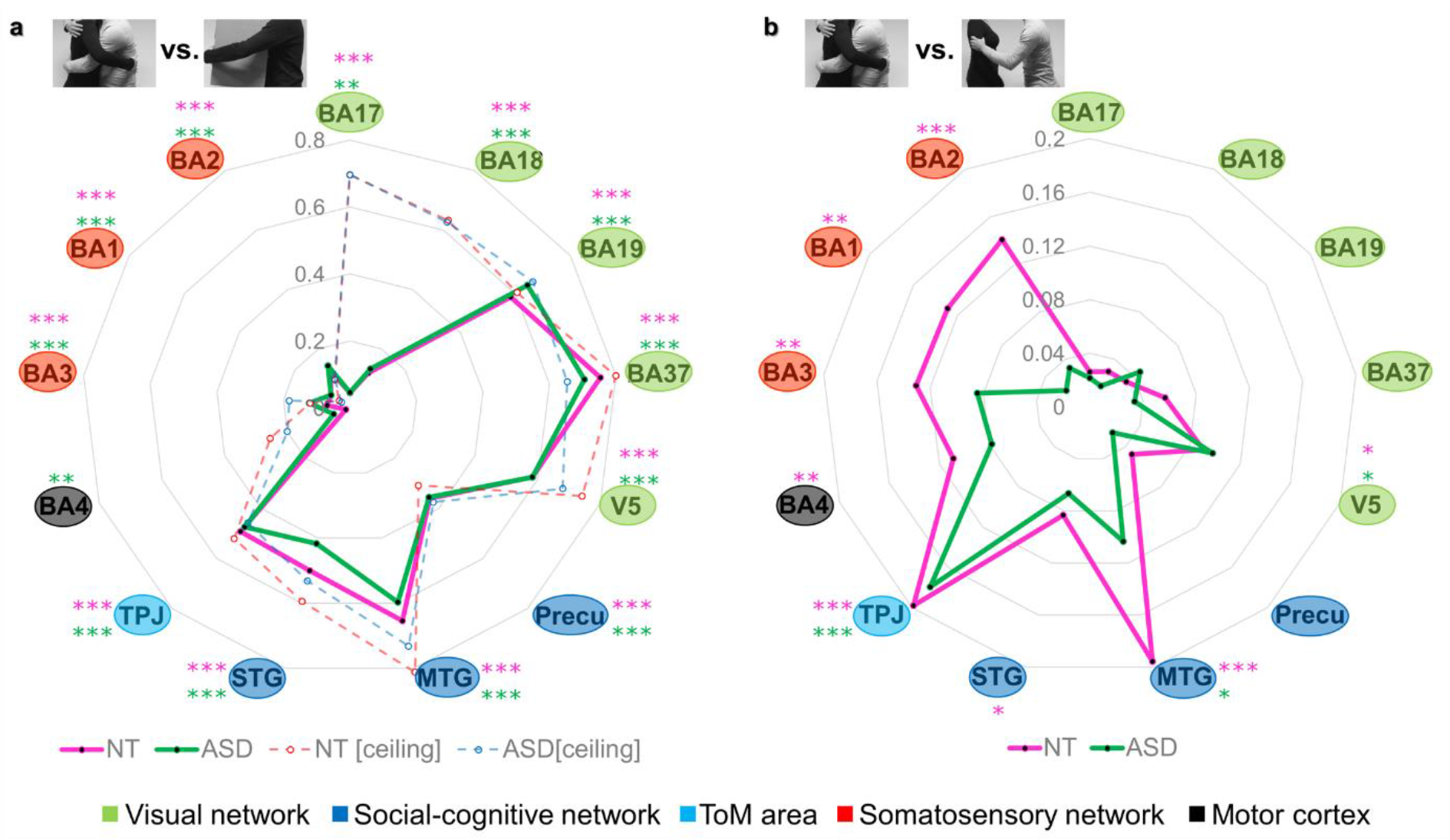
Neural representations of the social versus nonsocial distinction (a) and of affective meanings (b) in social touch scenes. Radar charts were used to plot the results of 13 ROIs in both groups, with a pink line used for the NT group and a green line used for the ASD group. Each of the 13 ROIs forms an individual axis that is radially arranged. The node (anchor) on the spoke (axis) represents the beta coefficient of each ROI. Figure (a) displays the beta coefficient from the multiple regression model in which the neural patterns of each of the ROIs were predicted based on the social vs. non-social factor. Figure (b) displays the beta coefficient from the multiple regression model predicting the neural patterns in each of the ROIs by means of perceived overall affect (i.e. the Euclidian combination of valence and arousal). The ROIs are ordered according to the brain network they belong to. The color of the circles surrounding the name of the ROI indicates the implied network (visual network in light green, social-cognitive network in blue, ToM area in sky blue, somatosensory network in red, and motor cortex in black). The single, double and triple asterisks indicate the statistical significance determined by the permutation tests at p < 0.05, p < 0.01, and p < 0.001 respectively in the NT group (pink asterisks) and the ASD group (green asterisks). In figure (a), we additionally plotted the correlation coefficient representing the noise ceiling of the neural data, which was derived from the reliability test (a red dashed line for the NT group and a blue dashed line for the ASD group).

### Neural representations underlying dimensions of overall affect in social touch observation

By combining the valence and arousal dimensions, we obtained a measure of overall affect conveyed in the social touch scenes. We investigated how this affective meaning of social touch is implemented in the brain when participants watch the interpersonal touch actions of others. In the NT group, statistically significant representations of overall affect were observed in V5 (β = 0.09, p = 0.04), MTG (β = 0.20, p < 0.001), STG (β = 0.08, p = 0.04), TPJ (β = 0.20, p < 0.001), the motor cortex (β = 0.11, p = 0.01), and the somatosensory cortex (BA3 β = 0.13, p = 0.002; BA1 β = 0.13, p = 0.004; BA2 β = 0.14, p < 0.001) (see the pink line in Figure 4b and Table S4). In the ASD group, however, significant representations of overall affect were found in V5 (β = 0.10, p = 0.05), MTG (β = 0.10, p = 0.03) and TPJ (β = 0.18, p < 0.001), but not in the somatosensory areas (BA3 β = 0.08, p = 0.08; BA1 β = 0.02, p = 0.32; BA2 β = 0.03, p = 0.32) (the green line in Fig. 4b and SI, Table S4).

Comparing both groups in terms of the strength of neural selectivity for the fine-grained socio-affective information in the core ToM area and somatosensory regions, we found equal representations in TPJ (t (40) = 1.04, p = 0.30), but significantly weaker representations in BA1 (t (40) = 3.06, p = 0.004) and BA2 (t (40) = 2.45, p = 0.02) in the ASD group. No significant group difference was found in BA3 (t (40) = 0.90, p = 0.37). The present results indicate that both groups are able to represent subtle socio-affective nuances of observed social touch interactions. However, whereas individuals with ASD only host this information in high-level visual areas (V5, MTG) and cognitively oriented ToM areas (i.e. TPJ), NT individuals additionally represent this information in a more embodied somatosensory format (BA3, BA1 and BA2). The neural representations of other factors, such as motor response and low-level visual features, were additionally examined. No group difference was found either in visual or in motor processing (SI, Fig. S4).

### Neural correlates of individual variability in touch avoidance and autistic traits

In our previous study in NT adults, we demonstrated that individual differences in the strength of neural representations of socio-affective touch in the somatosensory cortex were associated with individual differences in the attitude towards social touch in daily life [19]. Here we further extend these findings and connect the neuroscientific findings with core pervasive autistic traits.

### Social touch preference

When correlating the scores on the Social Touch Questionnaire (STQ) with the beta-coefficients indexing the quality of the overall affect representations in somatosensory cortex, the results indicated that a more positive attitude towards social touch is significantly associated with higher quality of overall affect representations in BA1 (rS = 0.43, p = 0.008) and BA2 (rS = 0.32, p = 0.04). Individual variability in affect representations in BA3 (rS = 0.08, p = 0.64) was not linked to the individual attitude towards social touch. Our results suggest that the functional organization and vicarious emotional sensitivity of the somatosensory cortex (i.e., BA1 and BA2) of individuals with a positive attitude towards receiving, witnessing, and providing social touch, may differ from the one of individuals who show social touch aversion.

### Social impairments

Likewise, the correlations between SRS-A scores and the strength of overall affect representations in somatosensory cortex revealed that individual differences in social responsiveness were significantly associated with the distinctness and specificity of overall affect representations in BA1 (rS = −0.38, p = 0.02) and BA2 (rS = −0.45, p = 0.003). Individual variability in affect representations in BA3 (rS = −0.12, p = 0.46) was not linked to individual differences in self-reported autistic traits.

Together, our results imply that the presence of affective representations of visually observed touch interactions in mid-to-high level somatosensory cortex (i.e., BA1 and BA2) shows an association with quantitative autism features and personal attitudes towards social touch. In particular, individuals who show higher social impairments tend to show a higher degree of social touch aversion, and both these characteristics are associated with less robust socio-affective representations in the mirror-somatosensory system during the observation of social touch.

## Discussion

Our study investigates the neural basis of socio-affective touch observation in adults with ASD compared to well-matched NT adults. In particular, we sought to clarify to what extent social impairments and touch aversion in ASD may be linked to aberrant cognitive representations of the affective aspects of touch (due to impaired ToM abilities) or to an inability to mobilize bodily representations (due to an absence of embodied simulation). Using fMRI-based RSA methods and a well-defined set of stimuli, we were able to pinpoint the atypicality of ASD in processing complex touch scenes containing multidimensional information (visual, somatosensory, and socio-affective). Our study provides novel evidence that adults with ASD specifically lack a differentiated somatosensory resonance when observing complex social touch interaction of others, despite a high degree of commonalities with NT adults in other aspects of neural information processing.

Adults with ASD rated the socio-affective touch scenes fairly similarly as NT adults, and both groups showed a high intra- and inter-subject consistency in their ratings. The only difference was in the perception of the positive touch scenes, such as a hug, which the adults with ASD rated slightly less pleasantly as compared to NT adults. While the effects are subtle, the results of this computer-based behavioral experiment are consistent with those of the questionnaire, which also revealed a significantly lower preference for receiving and observing social touch in daily life in adults with ASD. Similarly to Voos et al., [32], we also found an association between individual differences in social touch preference and the number of self-reported autism traits, while using different measurement instruments.

At the neural level, we found surprisingly high similarities between the two groups. Using both univariate and multivariate analysis approaches, intact neural selectivity for the social vs. non-social distinction of touch scenes was observed, in multiple brain regions including the ToM area and the somatosensory cortex. Although previous studies have shown that individuals with ASD may process social stimuli in an atypical manner [55], [56], the neural patterns associated with social vs. non-social touch scenes may still be distinctive as long as both conditions are perceived sufficiently differently from each other. This was indeed the case, also in the ASD sample, as illustrated by the ratings of valence and arousal in Fig. 1 (see SI for statistical comparisons between the two conditions). Accordingly, the current findings indicate that social impairments in ASD are not simply due to an inability to distinguish between social and non-social information.

Likewise, similar (univariate) neural activation in response to actual touch stimulation was observed in both groups in the current study. This observation contrasts with previous findings showing diminished neural response to affective touch events, especially to gentle brushstrokes, in individuals with ASD or in individuals scoring high on autistic traits [32], [57], [58]. Possibly, this discrepancy may be due to the administration of both positive and negative touch stimulation in our study, unlike the aforementioned studies which only delivered positive touch. Note that the current study aimed at investigating the neural representation of observed touch, and that the actual touch stimulation only served to confine a touch-related cortical area involving both positive and negative touch.

Strikingly, neural commonality with the NT group was even evident with regard to more delicate and fine-grained socio-affective processing. Individuals with ASD did represent subtle and differentiated information on the overall affect of socio-affective touch interactions in the TPJ, the most classical “social cognition ToM module”, suggesting intact rule-based mental state reasoning. For decades, impaired ToM ability has been put forward as the primary cause of socio-communicative impairments in ASD [7], [39], [43]. However, several recent studies have shown that high functioning and verbally gifted individuals with ASD, like the ones in our sample, successfully pass ToM tasks, possibly by using compensatory strategies [47]–[49].

Despite the typical ToM involvement in our ASD sample and despite the numerous behavioral and neural commonalities among both groups, we did observe significant and very specific differences in the more automatic and spontaneous processing of socio-affective touch interactions. Unlike NT adults, the ASD group did not show affective touch representations in mid-to-high level somatosensory areas (i.e., BA1 and BA2), indicating a lack of embodied resonance in relation to others’ bodily experiences. Our findings thereby extend studies demonstrating reduced empathic resonance to a painful touch experience of others as reflected in weaker mu suppression in ASD [31]. A lack of embodiment of others’ emotional state – not only painful sensations but also joyful ones – as revealed in the current study, further supports the argument that social difficulties in ASD may involve a lack of embodied simulation [59].

Lastly, building upon our previous study in NT adults [19], the current study provides evidence that individuals with stronger social touch avoidance or with more autistic traits experience diminished embodied somatosensory resonance with others. These findings extend recent studies that demonstrated an association between the level of activation in the somatosensory system during the observation of touch and inter-individual differences in empathy [23]–[26].

In sum, the current study provides strong support for the impaired embodied simulation account of ASD [40]–[42]. Accordingly, the less positive attitude towards reciprocal touch in ASD may be a consequence of the deficient automatic emotional resonance and the resulting increase in cognitive processing load during such interactions. Gallese and Sinigaglia [60] nicely illustrated the different formats of representations, and its impact, with the analogy of a route description: “Just as a map and a series of sentences might represent the same route with a different format, so might mental representations have partly overlapping contents while differing from one another in their format (e.g., bodily instead of propositional)” (p. 517). Crucially, while the same information can be represented in different formats, its utility is constrained by the format. Evidently, representing in a bodily format an emotion, such as disgust or pain, or a sensation, such as being touched, is different from representing them in a propositional format [60]. According to our findings and to the extent that TPJ may be a more cognitive and rule-based ToM module, individuals with ASD may not have access to the bodily format of the affective touch representations, but they have access to the propositional format. As a result, the depth of understanding and experiencing the state of others (and themselves) may differ between the two groups (i.e. “knowing” vs. “knowing and feeling”), which is also related to alexithymia in ASD [61]. The current findings may also motivate to reconsider how cognitive behavioral therapies designed to enhance mentalizing capacities of individuals with ASD can be complemented by more physical and bodily intervention strategies (e.g., mirror imitation therapy; [62], [63]) that directly target the deficient emotional resonance in clinical practice.

The current study also raises further questions which will require future research. In particular, the current study only included high functioning male adults with ASD. Although the homogeneity in our sample allowed controlling for confounding factors such as age, IQ, and gender, future studies may benefit from the inclusion of children and more severely affected individuals, including individuals with low IQ, who may not recruit compensatory cognitive strategies. Indeed, it remains an open question whether also these individuals would show intact rule-based ToM representations. Likewise, as the present study used a relatively effortless task, it remains to be seen whether intact ToM processing would still be in place when a task requires more higher-level cognitive exertion (e.g., understanding the meaning of touch based on the social norm and culture) [10].

## Materials and Methods

### Participants

21 male adults with a multidisciplinary ASD diagnosis (mean age = 25 years, standard deviation (SD) = 4.4, 3 left-handed) and 21 age-, gender-, and IQ-matched NT adults (mean age = 23.9, SD = 2.8, 4 left-handed) participated in the study. Detailed information on participants and recruitment procedure is described in SI. All participants provided written informed consent and were compensated for participating in this study approved by the KU Leuven Medical Ethical Committee (S59577).

### Stimuli

We used a recently created and well-validated set of 75 greyscale video clips (3 seconds each) displaying pleasant and unpleasant interpersonal touches (e.g., hugging, holding hands, slapping, caressing) and object manipulations (e.g., carrying a box). The 39 scenes for interpersonal or “social touch” and the 36 scenes for object manipulation or “non-social touch” were closely matched according to the type of interactions. For example, the movements involved in hugging another person versus holding a large box were matched. Various physical parameters from the video sequences were quantified, including pixel-wise intensity, pixel-wise motion energy, and total motion energy [4]. In the current study, the resulting parameters were defined as nuisance covariates in the multiple regression model.

Video clips and a detailed description of stimuli can be found in our previous study [4] (the link to the videos https://osf.io/8j74m/). Psychophysics Toolbox Version 3.0.12 (PTB-3) [64] in Matlab (R2015a, The Mathworks, Natick, MA) was used for stimulus presentation in all experiments.

### Behavioral rating of valence and arousal

First, participants took part in a behavioral experiment where they viewed all the video clips and reported the subjective feelings of pleasantness (“How pleasant is the touch?” 1-extremely unpleasant, 5-neutral, 9-extremely pleasant) and arousal (“How arousing is the touch?” 1-extremely calm to 9-extremely exciting) of 75 touch scenes. Each of the 75 stimuli was presented twice, with a short break in between the two sessions. More details about this experiment can be found in Experiment 2 of the previous study [4].

### Questionnaires

Next, participants filled out two questionnaires. The *Social Touch Questionnaire* (STQ) assesses individual attitudes towards receiving, offering, and witnessing social touch [65]. The *Social Responsiveness Scale for Adults* (SRS-A) is a normed self-report questionnaire measuring a wide range of behaviors characteristic of ASD [66]. Both STQ and SRS-A questionnaires are described in detail in SI.

### MRI acquisition

All participants underwent an MRI scanning session consisting of two functional MRI experiments and an anatomical scan. MRI images were acquired on a 3T Philips scanner with a 32-channel coil at the University Hospitals Leuven. MRI data acquisition parameters are described in detail in SI.

### fMRI Experiment 1: Receiving touch [A localizer run]

This experiment was used to localize the (affective) touch-related cortical areas as ROIs within the somatosensory cortex. Participants received pleasant (i.e. brush-stroke) and unpleasant (i.e. rubber band snapping) touch stimulations on the ventral surface of the right and left forearms while lying in the scanner. More information can be found in our previous study [19].

### fMRI Experiment 2: Observing touch [Main runs]

In the scanner, participants watched the same videos shown during the behavioral experiment while performing an orthogonal attention task (i.e., detecting the color of the shirt of the agent who initiates the touch, see SI for more information about the task). The main experiment consisted of 7 runs. In each run, the 75 videos were displayed in an optimally designed pseudorandom order in an event-related design. Each trial consists of a video presentation (3sc) and an inter stimulus interval (3sc) where a participant pressed a button to perform the task. More information about the task can be found in SI.

### Statistical analysis

Statistical inferences were made with appropriate tests (a parametric (e.g., two-tailed one sample t-test, two sample t-test, and Pearson correlation) vs. non-parametric test (e.g., Wilcoxon signed-rank test, Mann-Whitney U test, and Spearman correlation)) depending on the results of the Shapiro-Wilk normality test (with α < 0.05). For multiple regression analyses, we used a permutation test (details are described below).

### Behavioral data

The ratings of valence and arousal obtained through the two repetitions were averaged for each participant and stimulus. Then, the median of each group on perceived valence and arousal for each stimulus was determined. The ratings of the videos with positive, neutral, and negative emotional valence were analyzed separately. Based on the group valence rating, we assessed whether the participants had perceived positive touch scenes pleasantly, negative touch scenes unpleasantly, and non-social touch scenes neutrally. We compared the arousal ratings of social touch scenes with those of non-social touch scenes. A comparison of resulted ratings for positive, negative and non-social touch scenes was made between NT and ASD groups. Lastly, we performed within- and between-subjects reliability tests to examine how consistent the ratings are within and between participants in each group (SI).

In order to use the behavioral data as an independent variable to predict the neural data, we generated an overall affect score integrating the valence and arousal ratings. This was done by calculating the Euclidean Distance of valence and arousal ratings for each pair of videos, which was first done for each individual and then averaged across individuals. This operation resulted in an affective dissimilarity (distance) matrix. Note that there was a high within- and between-subject consistency of the behavioral valence and arousal ratings (SI), justifying the use of a group average affective dissimilarity matrix.

### Functional MRI data analysis

#### Preprocessing, first- and second-level analysis

Imaging data was processed using Statistical Parametric Mapping software (SPM 12). The standard preprocessing, first- and second-level analysis were implemented. Analysis pipelines are described in detail in SI.

During the preprocessing phase, we also assessed head movement of each participant and compared the two groups by using an Artifact Detection and Repair toolbox that calculates a composite measure of scan-to-scan movement. Runs whose maximum frame-wise displacement was greater than the voxel size (3 mm) were discarded (ASD = one run each from two participants; NT = one run from one participant). We found no group difference in the maximum (ASD = 1.38 mm, NT = 1.36, t(40) = 0.04, p = 0.97) and mean frame-wise head motion displacement (ASD = 0.13 mm, NT = 0.13, t(40) = 0.04, p = 0.96).

#### Regions of interest (ROIs)

We included the same ROIs as in our previous study [19]: Brodmann area (BA) 3, BA1, BA2, Parietal Operculum (PO), Insula, Middle Cingulate Cortex (MCC), Middle Temporal Gyrus (MTG), Superior Temporal Gyrus (STG), TPJ, Precuneus, BA17, BA18, BA19, BA37, V5, and BA4. All these ROIs are known to be involved in the processing of visually presented social touch scenes: vicarious touch processing in the somatosensory network (BA3, BA1, BA2, and PO, [67]), the pain network (insula and MCC, [68]), the social-cognitive network (MTG, STG, TPJ, and Precuneus, [11]), and the visual network (BA17, BA18, BA19, BA37, and V5, [69]). The motor cortex (BA4) was also included as the orthogonal task in the scanner involved active button presses, and thus motor responses.

We defined subject-specific ROIs by applying an identical procedure as employed in our previous study [19], including selecting the activated voxels within the anatomical mask for each ROI and trimming the overlapping voxels among the nearby ROIs. When the number of selected voxels was less than 10 per ROI, a more liberal threshold of P _uncorrected_ < 0.01 instead of P _uncorrected_ < 0.001 was used. Nevertheless, 9 out of 42 participants showed no activation in the insula and 11 out of 42 participants showed no activation in MCC. These two ROIs were therefore not included in the present study. This is not too surprising, given the low reliability estimates and limited explaining power of these ROIs in our previous study (cf. Fig. 5 and Fig. 7B in our previous study [19]). Exceptionally, BA4 was defined based on the anatomical mask only. We did not find any group differences in size of the ROIs (all *p* _uncorrected_ > 0.06). Mean ROI sizes and the p-values for individual ROIs are reported in SI. The ROIs are shown in Fig. 5.

**Fig. 5.**
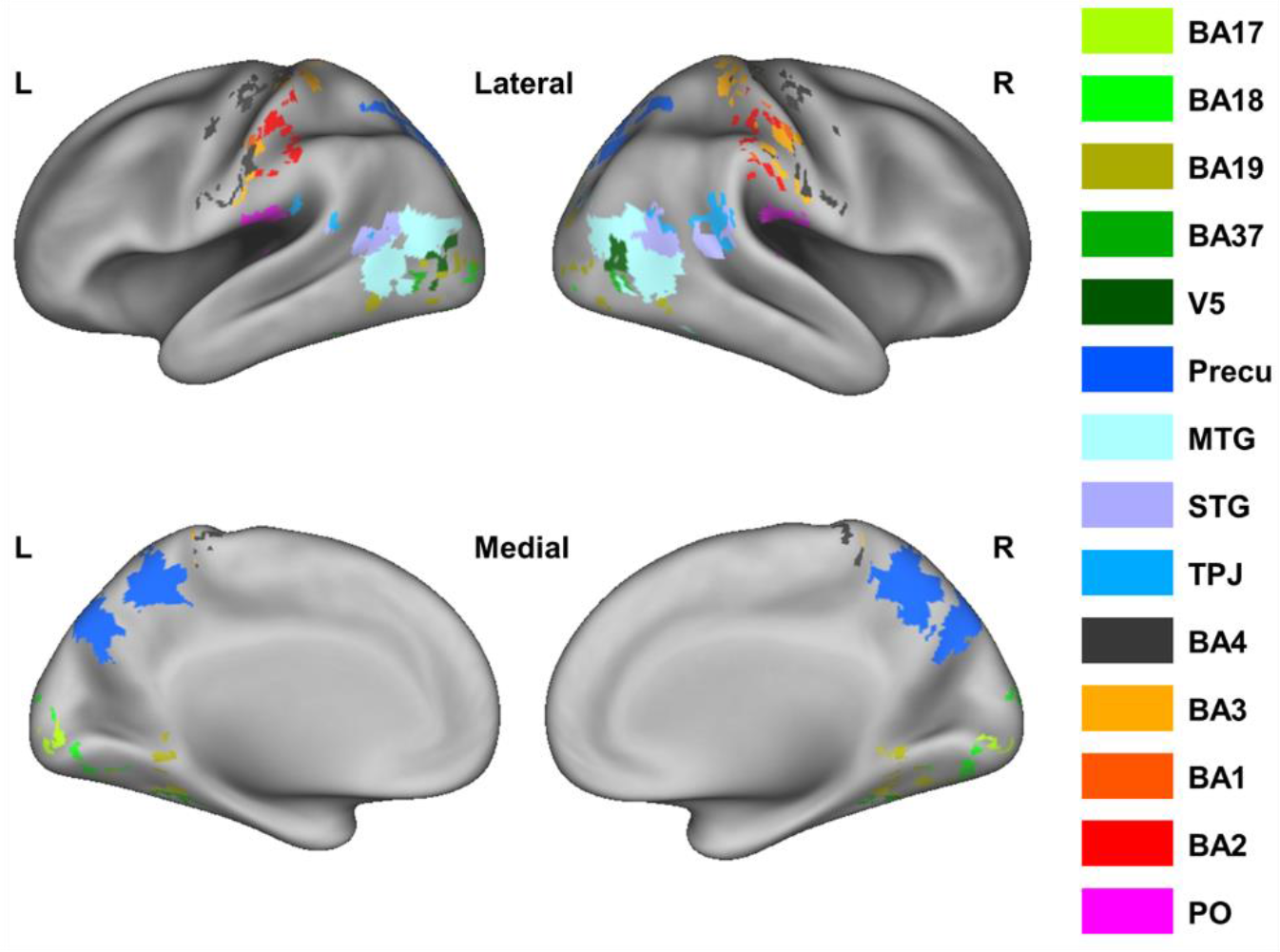
Visualization of ROIs. The figure illustrates the functionally-defined ROIs (except anatomically defined BA4), mapped on inflated cortices using the CARET software [70] with PALS atlas [71]. Note that mapping the volume-based data to surface can introduce artefacts.

#### Neural representational dissimilarity matrices (RDMs)

These matrices capture the difference in multi-voxel neural response patterns between pairs of videos. For example, if an ROI shows selectivity for the affective valence of the touch scenes, the neural patterns of two differing social touch scenes (e.g., hugging a person vs. slapping a person) will be largely dissimilar. On the other hand, if an ROI does not show this selectivity, the neural patterns will be largely similar across both types of affective interactions.

The “general touch RDM” involved the neural responses for both the social and nonsocial touch videos, and consisted of the pair-wise correlation coefficients of the 75 neural patterns. The “social touch RDM” exclusively involved the neural responses for the social touch videos, and consisted of the pair-wise correlation coefficients of the 39 neural patterns. We created these two RDMs per ROI and per participant and tested their reliability by applying an identical procedure as employed in our previous study [19]. The summarized procedure of making RDMs and performing a reliability test can be found in SI.

Note that based on the results of the reliability test, we excluded PO from further analysis as the between-subject variability was too high to conduct group analyses. In total, 42 (21 participants x 2 groups) general touch RDMs with 75 x 75 elements and 42 social touch RDMs with 39 x 39 elements were created per ROI. For the group analysis, we calculated the average of 21 individual RDMs to create each group’s general touch RDM and social touch RDM for each ROI. The RDMs of each ROI were used as dependent variables in each regression model in subsequent analysis.

#### Multiple regression analysis

To investigate which ROIs in which individuals host specific information on the displayed touch scenes, we carried out a series of multiple regression analyses to determine the independent contributions (as represented by the beta coefficients) of each variable of interest to the prediction of the neural data. Prior to this, we first vectorized each matrix, took only the upper-diagonal elements, and normalized the vector with Z-score transformation.

In the regression model predicting each group’s general touch RDM, the regressor variables consisted of a binary model of social versus non-social touch, the motor response made during the task, various physical parameters from the video sequences (pixel-wise intensity, pixel-wise motion energy and total motion energy), and the type of touch action.

In the regression model predicting each group’s social touch RDM, we replaced the binary model of social versus non-social touch by each group’s average affective evaluation of social touch (i.e., the affective dissimilarity matrix, see above). We used each group’s average affective dissimilarity matrix in each group’s regression model.

Statistical inferences for each group’s result were based upon a permutation test (1000 iterations), using the same procedure as described in [19]. We randomly shuffled the indices of the vector of neural data and computed the beta coefficients of each independent variable in a multiple regression model applied to the permuted data. We counted the number of times a beta coefficient obtained through this operation was greater than or equal to the observed value in the nonpermuted data. The result of dividing this number by 1000 became the empirical p-value after being corrected for multiple comparisons with the false discovery rate (FDR).

In order to directly compare the two groups taking into account inter-subject variability, we performed multiple regression on the neural matrices of individual participants in each group and applied either a two-sample t-test or a Mann-Whitney U test to compare the two groups. Against the background of the ToM and embodied simulation accounts (see Introduction), we specifically questioned the group difference in the quality of these representations in the core ToM area (TPJ) and the somatosensory cortex (BA3, BA1, and BA2), respectively.

## Acknowledgments

The authors thank all the participants who take part in this study. We also thank the clinical team of the Autism Expertise Center of UPC KU Leuven (Jean Steyaert, Stefaan Vertommen, Indra Struyven) for assistance with patient recruitment. This work was supported by a grant of the Marguerite-Marie Delacroix foundation to H.L.M (GV/B-336), and by grants of the Research Foundation Flanders (G088216N) and Excellence of Science program (G0E8718N; HUMVISCAT) to H.O.d.B. and B.B.

## Author contributions

H.L.M., H.O.d.B., and B.B. designed research. H.L.M., I.P., S.A., and S.V.D.P. performed research. S.A., and M.H. recruited the participants with autism. H.L.M analyzed data. H.L.M., H.O.d.B., and B.B. wrote the paper.

