## Supplemental Information for "Social touch observation in adults with autism: intact neural representations of affective meaning but lack of embodied resonance"

##### Participants

Participants with ASD had been diagnosed by a multidisciplinary team following DSM-IV or DSM-V criteria. None of the participants with ASD had neurological, psychiatric, or comorbid genetic conditions, such as epilepsy, traumatic brain injury, and attention deficit/hyperactivity disorder. They were recruited through the Leuven Autism Expertise Centre and the Psychiatric Clinic at the University Hospitals Leuven. On the other hand, adults with no prior diagnosis of ASD were recruited as NT participants through online advertising. None of the NT participants, nor first degree relatives, had a history of neurological, psychiatric, or medical conditions known to affect brain structure or function. None of the participants in both groups took psychotropic medication and all had normal or corrected-to-normal vision. Both groups were matched for gender (all male), age (ASD mean age = 25 years, SD = 4.4; NT mean age = 23.9, SD = 2.8; $t(40) = -1.0$, $p = 0.32$), and IQ (ASD mean IQ = 111.3, SD = 14.5; NT mean IQ = 111.5, SD = 12.3; $t(40) = 0.31$, $p = 0.76$), as assessed with the Wechsler Adult Intelligence Scale ((WAIS-IV-NL, [1]). The ASD participants generally show above average intelligence and show a fairly high level of social adaptive functioning (e.g., most have a regular job). There is a partial overlap (5 out of 21 NT participants) between the current NT sample and the data reported in [2]. These five participants were the only ones from the earlier study for which we had IQ scores and that are male.

##### Questionnaires

The *Social Touch Questionnaire* (STQ) comprises 20 items (e.g. “I generally like it when people express their affection towards me in a physical way”), and participants were asked to respond to each statement on a five-point scale (1 = strongly disagree, 2 = disagree, 3 = undecided, 4 = agree, 5 = strongly agree). A higher total score indicated that an individual expressed a stronger preference for reciprocal touch.

The *Social Responsiveness Scale for Adults* (SRS-A) comprises 64 items covering subscales for social communication and interaction and for restricted and repetitive patterns of behavior and interests. The SRS-A consists of three subscales measuring social deficits and one measuring restricted and repetitive behavior. A higher total score indicates a higher presence of quantitative autism traits.

#### MRI data acquisition

Functional imaging was performed with a gapless, echo planar imaging sequence (repetition time (TR) = 2000 ms, echo time (TE) = 30 ms, flip angle (FA) = 90°, field of view (FOV) = 216 × 216 mm, in-plane matrix = 80 × 80, voxel size = 2.7 x 2.7 × 3 mm, 37 slices), with the acquisition of 239 volumes for each run of the main experiment and 298 volumes for the localizer run.

Structural MR images were collected using a T1-weighted sagittal high-resolution magnetization prepared rapid gradient echo (MPRAGE) sequence [TR = 9.6 ms, TE = 4.6 ms, FA = 8°, FOV = 250 × 250 mm, in-plane matrix = 256 × 256, voxel size = 0.98 × 0.98 × 1.2 mm, 182 axial slices].

#### MRI data preprocessing

Preprocessing steps include 1) slice timing correction, 2) realignment of functional images to the mean image of the first run (the images from the localizer experiments were also aligned to this mean image), 3) registration of the anatomical image to the functional images, 4) segmentation which provides forward deformation fields, 5) normalization process which warps all structural and functional images to a Montreal Neurological Institute (MNI) space with a re-sampling size of 3 x 3 × 3 mm based on forward deformation fields, and 6) spatial smoothing. Gaussian kernels with a 5 mm full-width at half maxima (FWHM)) was chosen as the smoothing parameter, except for the second-level univariate group analysis (8 mm FWHM).

#### Main fMRI Experiment: Observing touch and the task

Participants were instructed to indicate with a button press during the 3s inter stimulus interval (ISI) whether the person in the video clip who initiated the touch was wearing a black sweatshirts (left hand button) or a grey sweatshirts (right hand button). All the videos were projected on a screen behind the scanner and participants viewed them through a mirror mounted on the head coil.

Every run consisted of 3 blocks each of which contained a baseline condition displaying a fixation cross (6sc) and 25 trials consisting of video presentation (3sc) and ISI (3sc). The total duration of each run took 7.8 min [3 blocks x ((baseline (6sc) + 25 trials x (video presentation (3sc) + ISI (3sc))].

#### First- and second-level analysis

The first-level (subject-level) analysis was carried out with a standard general linear model (GLM). Delta functions were used to model the regressors for the main runs (observing touch) while boxcar functions were used for the localizer run (receiving touch). The regressors of interest differed across four GLMs.

1) The first GLM was created for a second-level univariate group analysis and was fitted to the 8 mm FWHM smoothed preprocessed imaging data. This GLM contained three predictors, i.e. social, non-social, and baseline condition, referring to the social touch videos, the non-social touch videos, and a fixation cross, respectively. The resulting data were used for standard random-effect group-level whole brain analyses were conducted. We identified significantly activated voxels in the "social minus non-social touch observation" contrast, and vice versa, and we compared brain activation between the two groups for each contrast. All statistical maps were thresholded at $P_{FWE} < 0.05$.

2) The second GLM was created for a representational similarity analysis (RSA) and was fitted to the 5 mm FWHM smoothed preprocessed imaging data. This GLM contained 75 predictors, i.e., one regressor for each video stimulus. We used the resulting 75 estimated beta-values as input for MVPA to construct a subject-specific neural dissimilarity matrix (see section 8).

Two more GLMs were additionally fitted for defining ROIs. 3) The third GLM (fitted to the 5 mm FWHM smoothed data from the main experiment) again contained the three predictors social touch observation, non-social touch observation, and baseline (fixation cross). We used the contrast of all the touch videos versus baseline to identify the majority of ROIs, except for the touch-related ROIs. 4) The last GLM (fitted to the 5 mm FWHM smoothed data from the localizer experiment) contained three predictors: pleasant touch, unpleasant touch, and rest condition. We used the contrast of both touch conditions versus rest to identify the first-hand touch-selective cortical regions as ROIs.

#### Mean ROI sizes

Mean ROI sizes after trimming and p-values for group comparisons were as follows: BA3 = 81 (ASD = 78; TD = 83; p = 0.67); BA1 = 21 (ASD = 19; TD = 23; p = 0.17); BA2 = 67 (ASD = 62; TD = 71; p = 0.38); PO = 109 (ASD = 97; TD = 120; p = 0.21); Precuneus = 447 (ASD = 414; TD = 480; p = 0.44); MTG = 404 (ASD = 355; TD = 453; p = 0.06); STG = 197 (ASD =

176; TD = 219; p = 0.26); TPJ = 116 (ASD = 104; TD = 127; p = 0.43); BA17 = 88 (ASD = 86; TD = 90; p = 0.57); BA18 = 290 (ASD = 283; TD = 296; p = 0.66); BA19 = 247 (ASD = 233; TD = 260; p = 0.29); BA37 = 94 (ASD = 82; TD = 107; p = 0.06); V5 = 42 (ASD = 41; TD = 43; p = 0.50); BA4 = 386 voxels. Here, reported p-values were not adjusted for multiple comparisons corrections. The p-values corrected for the false discovery rate (FDR) for multiple comparisons are all above 0.42.

#### Neural representational dissimilarity matrices (RDMs)

The procedure of constructing RDMs include the following steps: 1) extracting beta-values of all voxels in each ROI for each stimulus, 2) per run normalizing the beta-values for each voxel in each ROI by subtracting the average beta-value over all conditions, 3) generating multi-voxel patterns for each stimulus by averaging the normalized beta-values of seven runs, 4) generating two symmetric matrices (i.e., 75 x 75 general touch matrix and 39 x 39 social touch matrix) per ROI by computing the Pearson correlation coefficients between multi-voxel patterns of each possible pair combination of stimuli (RSA, [3]), 5) lastly, transforming the matrix into RDM by subtracting correlation coefficients from 1.

#### Reliability test

Two reliability tests were performed – one for each participant group separately – to test whether an RDM of each ROI contains a reliable signal.

First, by comparing the diagonal of RDMs (diagonal = comparison of a condition with itself) with the non-diagonal, we verified whether multi-voxel patterns of a particular ROI are worth further analysis [4]. The procedure included: 1) randomly splitting the seven runs into two halves, 2) calculating the average matrix of each, 3) correlating the two, 4) placing the resulting correlation coefficients of the within-conditions and the between-condition correlations on the diagonal and non-diagonal elements respectively of a 75 × 75 square matrix, 5) repeating the aforementioned steps for 100 times, and 6) calculating the average of 100 square matrices per ROI and group. To measure whether the multi-voxel patterns contain a meaningful signal, we compared the correlation coefficients of the diagonal cells of the matrix created with the aforementioned methods with those of the shuffled matrix of which indices were randomly rearranged. Statistical inferences were based upon a permutation test (1000 times of iterations). We counted the number of times a shuffled matrix contained an averaged diagonal value that was greater than or equal to that of an original matrix and then divided the number by 1000.

The result of dividing this number by 1000 became the empirical p-value after being corrected using the FDR for multiple comparisons (cf. the number of ROIs). Based on this analysis, for both groups, we did not exclude any ROI from further analysis.

Second, with the split-half between-subject correlational method [5], we estimated the maximum regression coefficient we could expect in predicting the neural data with other variables while taking into account between-subject variability in the multi-voxel patterns. 21 RDMs of each group were randomly split into two halves and averaged. We vectorized each matrix, took only the upper-diagonal elements, and correlated a resulting vector of one matrix with that of another matrix. Finally, we adjusted the resulting correlations with the Spearman-Brown formula ($2 \times r/(1 + r)$). We repeated the aforementioned steps 100 times, each time randomly designating participants into two sub-groups. In the end, the results are averaged per ROI and group. P-values were corrected using the FDR for multiple comparisons (cf. the number of ROIs). Based on the results, we excluded PO (no correlation between participants, $r = 0.009$) from the further analysis.

### Results

#### Affective responses to social and non-social touch videos

Both groups perceived positive touch videos (e.g., hugging a person) as pleasant (NT, median = 7.4 (the median absolute deviation (MAD) = 0.4); ASD, median = 6.8 (0.6)), whereas the negative touch videos (e.g., slapping a person) were rated as unpleasant (NT, median = 2.9 (0.3); ASD, median = 3 (0.5). Non-social touch videos (e.g., carrying a box) were rated as neutral (NT, median = 4.8 (0.2); ASD, median = 4.8 (0.2). A Wilcoxon signed-rank test revealed that rated valence was significantly higher for the positive touch videos than the negative ones in both groups (NT, $z = 4.02$, $p < 0.001$; ASD, $z = 3.98$, $p < 0.001$). In line with this, in both groups, we observed significant differences in the rated valence between positive videos and non-social ones (NT, $z = 4.01$, $p < 0.001$; ASD, $z = 3.29$, $p < 0.001$) and negative ones and non-social ones (NT, $z = - 4.02$, $p < 0.001$; ASD, $z = - 3.98$, $p < 0.001$).

Concerning arousal ratings, both groups perceived social touch as exciting (NT, median = 5.7 (0.9); ASD, median = 6.2 (0.8)) and non-social touch stimuli as calm (NT, median = 2.4 (0.9); ASD, median = 2.5 (0.6)). The rated arousal was significantly higher for the social touch videos than the non-social ones in both groups (NT, $z = 4.01$, $p < 0.001$; ASD, $z = 4.01$, $p < 0.001$).

#### Intra- and inter-subject consistency of valence and arousal ratings

We conducted within- and between-subjects reliability tests on resulting ratings for each group. For the within-subject reliability tests, we calculated the Spearman correlation for the ratings of 75 stimuli between the two test sessions for each participant. For the between-subjects reliability test, a split-half correlation method was used, measuring the correlation between the ratings of one half of the participants and those of the other half of the participants.

The results revealed that participants were consistent in their ratings between the two sessions (NT, valence median $r_S = 0.90$, $p < 0.001$; arousal median $r_S = 0.85$, $p < 0.001$; ASD, valence median $r_S = 0.85$, $p < 0.001$; arousal median $r_S = 0.77$, $p < 0.001$). Figure S1a illustrates data points of individual participants' correlations between the two sessions for valence ratings. A Mann-Whitney U test revealed that two groups show no difference in within-subject consistency for valence ($z = 0.98$, $p = 0.33$) and arousal ratings ($z = 1.43$, $p = 0.15$), indicating that individuals with ASD rated the valence and the arousal of the stimuli as coherently as NT adults across two sessions.

Concerning inter-subject consistency, the results revealed high consistency for valence (NT, median $r_S = 0.85$, $p < 0.001$; ASD, median $r_S = 0.74$, $p < 0.001$) and arousal ratings (NT, median $r_S = 0.73$, $p < 0.001$; ASD, median $r_S = 0.69$, $p < 0.001$) within the group (Figure S1b). Unlike intra-subject consistency, however, significant differences in inter-subject consistency for the valence ($z = 9.58$, $p < 0.001$) and arousal ratings ($z = 2.32$, $p = 0.02$) were observed, suggesting more heterogeneity of ratings in the ASD group despite the high consistency.

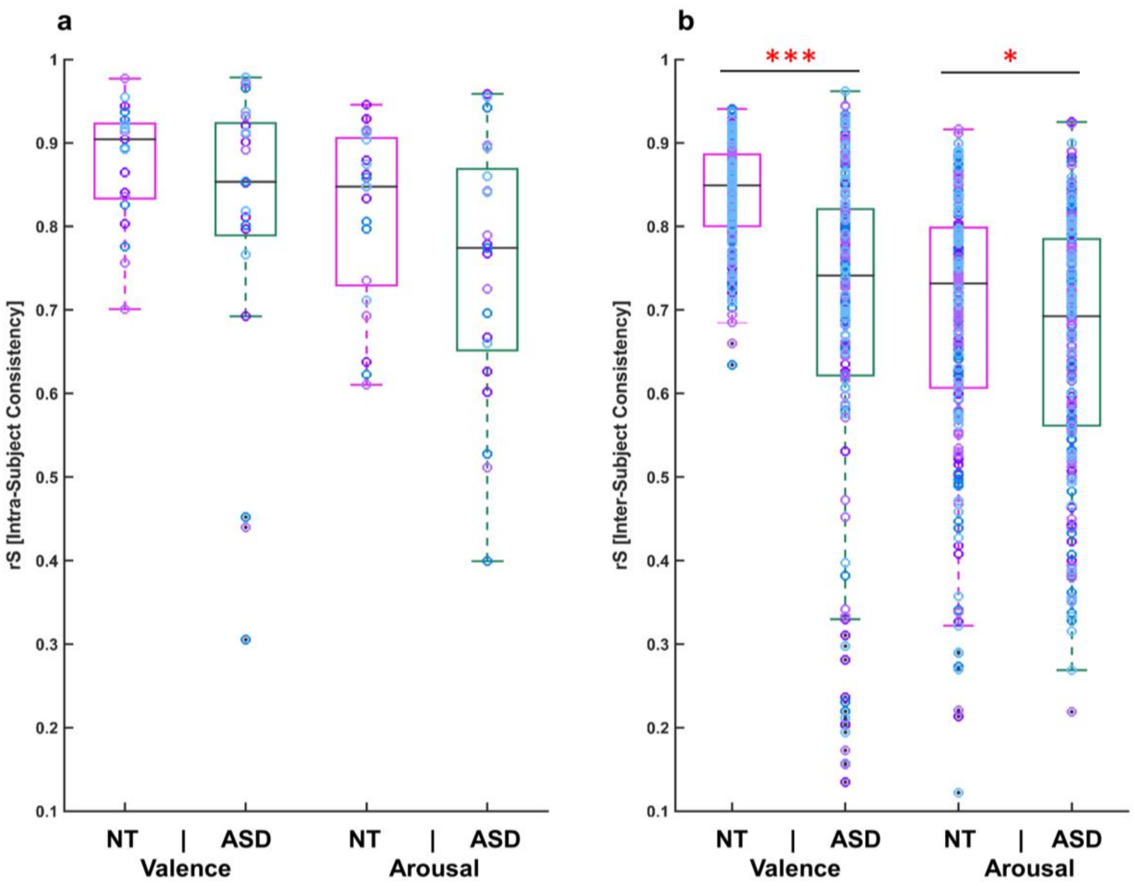


**Fig. S1. Intra-subject, and inter-subject consistency.** The figure illustrates intra- (a) and inter-subject consistency (b) on valence and arousal ratings in both groups. The central black lines inside of the boxes indicate the group medians of Spearman correlations. The bottom and top border edges of the boxes illustrate the 25$^{th}$ and 75$^{th}$ percentiles respectively; the whiskers illustrate the range of the rank correlation coefficients within 99.3 % coverage. All data points are additionally marked as purple/sky-blue circles. The black dots dawn outside of the whiskers indicate the outliers. The single red asterisk indicates the statistical significance at $p < 0.05$ and three red asterisks show statistical significance at $p < 0.001$.

**Table S1 The strength and direction of a linear relationship between the social touch behavior and three subscale scores of SRS-A.**

| | **Social Awareness** | **Social Communication** | **Social Motivation** |
|---|---|---|---|
| **NT** | r = -0.42, p = 0.06 | **r = -0.54, p = 0.01** | **r = -0.67, p < 0.001** |
| **ASD** | r = -0.4, p = 0.08 | **r = -0.59, p = 0.005** | r = -0.36, p = 0.11 |

Bold text indicates the significant association between the two variables.

### Neural responses to observed and felt touch

#### Observing touch

A second-level univariate analysis across all participants revealed that the current results of the contrast between social and non-social touch observation conform well to our previous findings [2]. Increased brain activity was observed in multiple brain areas, including limbic (i.e., the anterior insular cortex) and social brain regions (e.g., the superior temporal gyrus (STG), the TPJ, the medial prefrontal cortex (MPFC), and the precuneus) during the observation of social touch as compared to non-social touch (thresholded at $p_{FWE} < 0.05$). On the other hand, the fusiform gyrus (FuG), a high-level visual region implicated in object recognition, was relatively more active during the observation of non-social touch videos (involving a person handling an object).

**Table S2 Brain areas activated during the observation of social touch compared to non-social touch and vice versa.**

| *Social Touch Observation > Non-Social Touch Observation* | | | | | | |
|---|---|---|---|---|---|---|
| | Peak X | Peak Y | Peak Z | T(40) | P | N Voxels |
| *R MTG/IOG/AnG* | 48 | -58 | 11 | 20.5 | 0.000 | 3642 |
| *R STG/SMG/AnG* | 60 | -40 | 20 | 17.3 | 0.000 | / |
| *R IOG* | 48 | -73 | -1 | 15.7 | 0.000 | / |
| *R MFG/PreG* | 42 | 5 | 44 | 15.7 | 0.000 | 1194 |
| *R AIns/FO/IFG* | 33 | 26 | 2 | 12.1 | 0.000 | / |
| *R MFG/PreG/SFG* | 33 | -1 | 53 | 10.3 | 0.000 | / |
| *L AIns/FO/IFG* | -30 | 23 | 5 | 13.4 | 0.000 | 627 |
| *L PreG/MFG* | -39 | -4 | 53 | 10.5 | 0.000 | / |
| *L PreG/MFG* | -45 | 2 | 35 | 8.1 | 0.000 | / |
| *R SMC/SFG* | 9 | 14 | 53 | 13.2 | 0.000 | 539 |

| | Peak X | Peak Y | Peak Z | T(40) | P | N Voxels |
|---|---|---|---|---|---|---|
| *L SMC/MCgG* | -6 | 14 | 47 | 12.4 | 0.000 | / |
| *R SMC/SFG* | 9 | 5 | 62 | 8.05 | 0.000 | / |
| *R LiG* | 18 | -82 | -10 | 12.2 | 0.000 | 211 |
| *R LiG* | 12 | -88 | -4 | 9.7 | 0.000 | / |
| *R OCP* | 15 | -94 | 8 | 8.7 | 0.000 | / |
| *L OCP* | -21 | -94 | 11 | 8.7 | 0.000 | 62 |

***Non-Social Touch Observation > Social Touch Observation***

| | Peak X | Peak Y | Peak Z | T(40) | P | N Voxels |
|---|---|---|---|---|---|---|
| *L FuG/LiG* | -27 | -49 | -10 | 16.6 | 0.000 | 476 |
| *L FuG/Hippo/ParaHioppo* | -30 | -34 | -16 | 11.68 | 0.000 | / |
| *R LiG/FuG* | 24 | -46 | -10 | 15.8 | 0.000 | 433 |
| *R FuG/ Hippo/ParaHioppo* | 33 | -19 | -25 | 6.7 | 0.001 | / |

**Receiving touch**

As with the observed touch paradigm, the second-level univariate analysis revealed that the current results of felt touch minus rest condition conform well to our previous findings [2]. Increased brain activity was observed in multiple brain areas including limbic (i.e., the posterior insular cortex and the middle cingulate gyrus) and somatosensory areas (i.e., the postcentral gyrus and the parietal operculum) when receiving touch as compared to rest ($p_{FWE} < 0.001$) (Fig. S2. and Table S3). No group differences were found in this experiment, which in the current context was mainly used as a localizer to help define the somatosensory regions.

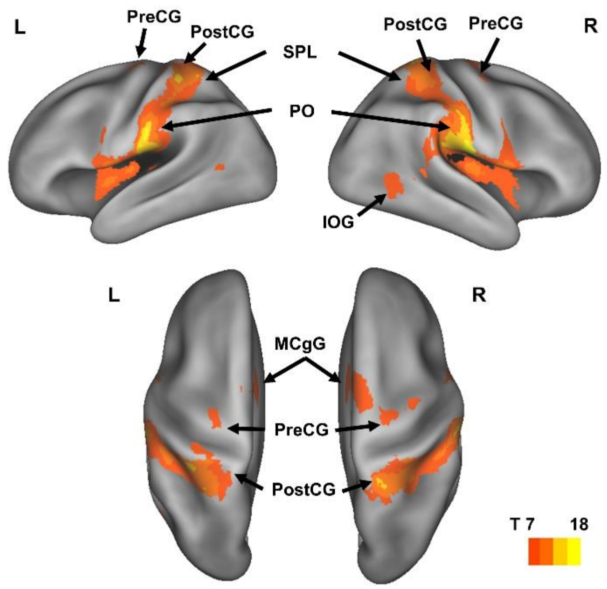


**Fig. S2. Brain areas showing increased neural activation for receiving touch.** The figure shows the mean group effects of felt touch minus rest (p FWE < 0.001, k= 60). We mapped the contrast result on inflated cortices with PALS atlas using CARET software. MCgG = middle cingulate gyrus, PO = parietal operculum.

**Table S3 Brain areas activated when receiving touch compared to resting.**

| ***Receiving Touch > Rest*** | | | | | | |
|---|---|---|---|---|---|---|
| | Peak X | Peak Y | Peak Z | T(40) | P | N Voxels |
| PO | 57 | -19 | 20 | 18.2 | 0.000 | 2778 |
| *SPL/PostG* | 24 | -43 | 68 | 14.2 | 0.000 | / |
| *SPL/PostG* | 51 | -25 | 35 | 13.2 | 0.000 | / |
| *PO* | -51 | -25 | 17 | 16.9 | 0.000 | 2293 |
| *PostG/SMG* | -57 | -22 | 29 | 16.4 | 0.000 | / |
| *PostG/SPL* | -36 | -37 | 56 | 13.6 | 0.000 | / |
| *Cerebellum* | 18 | -70 | -22 | 12.8 | 0.000 | 359 |
| *Cerebellum/FuG* | -27 | -55 | -25 | 12.5 | 0.000 | / |
| *Cerebellum/FuG* | 27 | -58 | -25 | 12.4 | 0.000 | / |

| | | | | | | |
|---|---|---|---|---|---|---|
| *SFG/SMG* | 9 | -1 | 71 | 10.4 | 0.000 | 806 |
| *SMG/MCgG* | 0 | -1 | 53 | 9.6 | 0.000 | / |
| *SFG/SMG* | -9 | -4 | 74 | 9.4 | 0.000 | / |
| *MTG/IOG/MOG* | -54 | -70 | 8 | 9.5 | 0.000 | 66 |

#### Neural representations underlying observed social versus non-social touch processing

Both NT and ASD participants showed high multi-voxel selectivity for the distinction between social and nonsocial touch scenes. The neural patterns of BA37 (NT $\beta = 0.75$, $p < 0.001$; ASD $\beta = 0.71$, $p < 0.001$) and of MTG (NT $\beta = 0.65$, $p < 0.001$; ASD $\beta = 0.60$, $p < 0.001$) were the best predicted by the social vs. non-social variable. Furthermore, in both groups, the social vs. non-social variable could explain changes in the distributed patterns of neural activity in the somatosensory regions (e.g., BA 2 NT $\beta = 0.13$, $p < 0.001$; ASD $\beta = 0.14$, $p < 0.001$), implying intact representations of the rough social vs. non-social subdivision of the observed touch scene in somatosensory regions of the ASD group. Likewise, in both groups, TPJ showed high neural selectivity for the social vs. non-social contrast. Table S4 and Fig. S3. contain a complete list of results.

Additionally, the results of between-subject reliability test revealed that the amount of variation in the neural data explained by the social vs. non-social model reached almost the noise ceiling in the areas BA37, MTG, TPJ, BA1, BA2 and BA3 (e.g., BA 37 of the NT group r (the ceiling) = 0.80 and $\beta = 0.75$, Fig. S3a and b). In contrast, the social vs. non-social model ($\beta = 0.05$) did not account for much of the neural patterns in the early visual cortex, although these areas exhibit high noise ceilings (e.g., BA17 of the NT group $r = 0.70$). It implies that other (more low-level) regressor variables may have a greater impact on explaining the neural data of these areas (Fig. S4b).

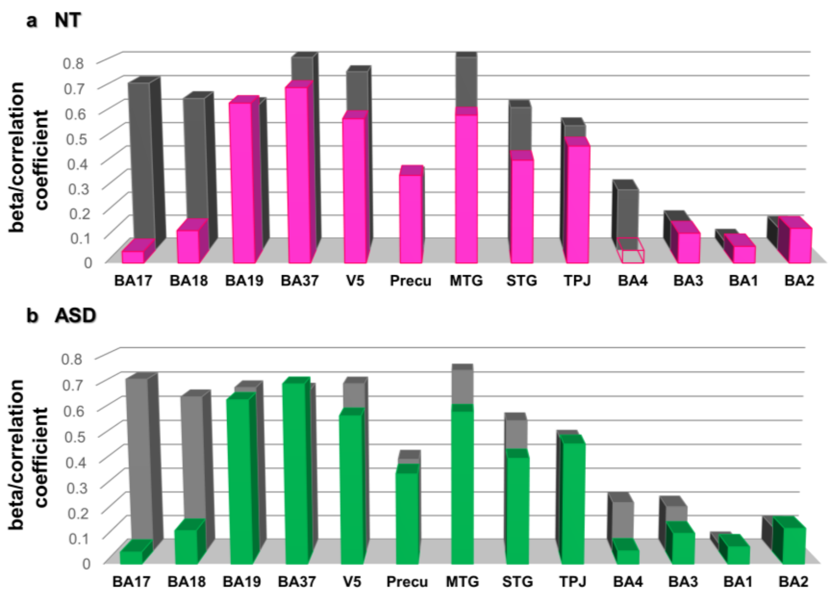


**Fig. S3 Representation of the social vs. non-social touch and reliability estimate in ROIs.** The Y-axis illustrates the predictor's estimated beta coefficient (social vs. non-social model) indicating the slope of the regressor in the multiple regression model predicting the neural patterns of each ROI in the NT (a, pink bars) and the ASD (b, green bars) groups. The grey bars in both figure (a) and (b)) depict the correlation coefficients representing a noise ceiling (X-axis lists the name of the ROIs. The bar without the color-fill (BA4 in figure a) means that the results are not significant.

**Table S4 The beta coefficients of the social/non-social and overall affective dimensions for all ROIs and both groups.**

| | *Social/Non-Social* | | *Affect* | |
|---|---|---|---|---|
| | NT | ASD | NT | ASD |
| *BA17* | 0.05** | 0.05** | 0.03 | 0.02 |
| *BA18* | 0.12*** | 0.13*** | 0.03 | 0.02 |
| *BA19* | 0.58*** | 0.64*** | 0.03 | 0.05 |
| *BA37* | 0.75*** | 0.71*** | 0.06 | 0.03 |
| *V5* | 0.58*** | 0.58*** | 0.09* | 0.10* |
| *Precuneus* | 0.36*** | 0.36*** | 0.05 | 0.03 |
| *MTG* | 0.65*** | 0.60*** | 0.20*** | 0.10* |

| | | | | |
|---|---|---|---|---|
| *STG* | 0.50*** | 0.42*** | 0.08* | 0.07 |
| *TPJ* | 0.49*** | 0.47*** | 0.20*** | 0.18*** |
| *BA4* | 0.01 | 0.05** | 0.11** | 0.08 |
| *BA3* | 0.07*** | 0.12*** | 0.13** | 0.08 |
| *BA1* | 0.07*** | 0.07*** | 0.13** | 0.02 |
| *BA2* | 0.13*** | 0.14*** | 0.14*** | 0.03 |

Asterisks denote FDR-corrected p values from the permutation test (* $p < 0.05$, ** $p < 0.01$, *** $p < 0.001$).

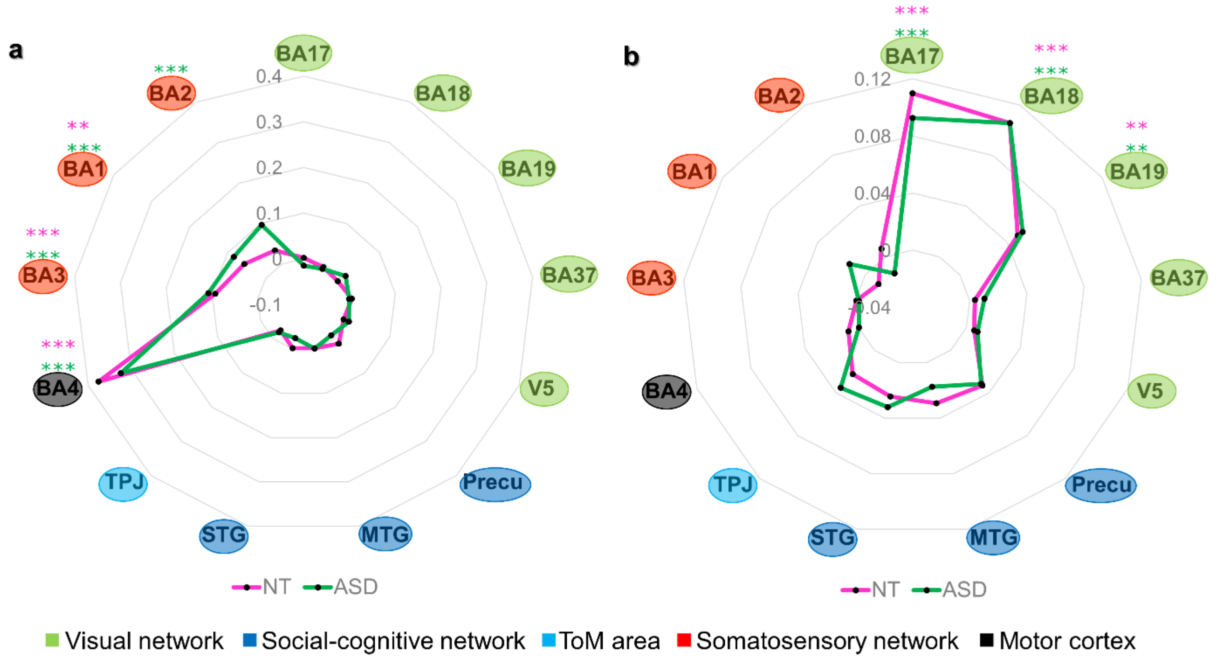


**Fig. S4. Neural representations of motor responses made during the task and the pixel-wise intensity of the video frames.** Radar charts were used to plot the results of 13 ROIs in both groups (a pink line used for the NT group and a green line used for the ASD group) on the same graph. Each of the 13 ROIs forms an individual axis that is radially arranged. The node (anchor) on the spoke (axis) represents the beta coefficient of each ROI. The figure (a) and (b) show the beta coefficient of each ROI from the multiple regression model in which the neural patterns of the ROIs were predicted based on motor reponses made during the task (a) and pixel-wise intensity of the videos (b) respectively. A circle filled with a color (a green circle used for the ASD group, and a purple circle used for both groups) surrounding the name of the ROI indicates the beta coefficient being statistically significant, which was driven from the permutation test. The figure indicates the motor and visual representations in the motor cortex and the occipital lobe respectively in both groups, highliting the intact motor and visual processing in ASD.
